# The IL-1 Family Controls Acute Mucosal Fungal Infection and Mucosal-Systemic Dissemination

**DOI:** 10.64898/2026.04.10.717440

**Authors:** James S. Griffiths, Alexander Kempf, Robert J. Pickering, Emily L. Priest, Olivia K. A. Paulin, Léa Lortal, Andrew Donkin, Olivia W. Hepworth, Don N. Wickramasinghe, Aize Pellon, Adrian Lau, Harun Papini, Sarah L. Gaffen, Jonathan P. Richardson, Julian R. Naglik

**Affiliations:** Centre for Host-Microbiome Interactions, Faculty of Dentistry, Oral C Craniofacial Sciences, King’s College London, United Kingdom; The Institute of Cancer Research, London, United Kingdom; Department of Biochemistry and Biophysics, University of California San Francisco, San Francisco, USA; Division of Infectious Diseases, Department of Medicine, Massachusetts General Hospital, Boston, Massachusetts, USA; Harvard Medical School, Boston, Massachusetts, USA; Inflammation and Macrophage Plasticity Laboratory, CIC bioGUNE-BRTA (Basque Research and Technology Alliance), Derio, Spain; Division of Rheumatology and Clinical Immunology, University of Pittsburgh, USA

**Author notes:** Contributed whilst at^1^ (Léa Lortal, Olivia W. Hepworth, Aize Pellon).

## Abstract

*Candida albicans* is a major opportunistic pathogen in humans that is capable of breaching mucosal barriers and causing severe systemic infections with high mortality. How the host controls mucosal infection and prevents dissemination remains unclear but is essential for improving disease outcomes. Here, we demonstrate that *C. albicans* induces specific IL-1 family members, which are critical for initiating mucosal protection by controlling antimicrobial peptides, IL-17, and neutrophil responses. Loss of combined IL-1 family signalling led to severe mucosal *C. albicans* infection, which was eventually resolved by a potent neutrophil response. However, in neutropenic conditions (a key risk patient factor) abolishing IL-1 family signalling resulted in *C. albicans* dissemination, predominantly to the liver, mirroring clinical disease and leading to mortality. This study highlights the IL-1 family as a key initiator of mucosal immunity, restricting mucosal invasion and cooperating with neutrophils to prevent life- threatening systemic infections.

## Introduction

Fungal pathogens are responsible for an increasingly severe global disease burden that kills ∼2.5 million individuals every year. The fungus *Candida albicans* (*C. albicans*) is a key contributor to this burden and is implicated in almost 1 million deaths annually [1]. *C. albicans* is typically a harmless constituent of a healthy microbiota but if left unchecked can overgrow its niche and drive superficial mucosal infection [2]. While mucosal infection is common and contributes to morbidity, it is invasive systemic disease that drives mortality. Systemic *C. albicans* infections arise from uncontrolled mucosal populations and from the colonisation of medical plastics, coupled with ineffective host immunity [3, 4]. Immunocompromised patients such as those with neutropenia (often a result of cancer treatment), graft versus host disease, and HIV are at risk of invasive systemic disease [4]. With increasing resistance to antifungals, poor diagnostic tools and limited therapeutics, understanding how *C. albicans* mucosal infections develop and, critically, how they disseminate, is vital to managing *C. albicans* disease.

*C. albicans* is a polymorphic yeast that can produce invasive hyphal filaments that infect mucosal tissues. *C. albicans* hyphae secrete candidalysin, a toxin that damages mucosal barriers and activates innate host immune responses [5]. Mucosal tissues respond to the presence of candidalysin by inducing signal transduction pathways that promote the release of anti-microbial peptides (AMPs), proinflammatory cytokines, IL- 17 production (a key protective barrier cytokine) and neutrophil recruitment [5–8]. In an immunocompetent host this response usually resolves pathogenic fungal overgrowth, but immunocompromised hosts often struggle to contain the infection and develop superficial candidiasis. Uncontrolled mucosal candidiasis can persist, penetrate through the mucosa resulting in invasive candidiasis, and then disseminate to organs including the liver and spleen. However, while invasive disease is a severe clinical complication with high mortality (10-77%) [3, 4, 9], not all susceptible patients develop invasive *C. albicans* infections, suggesting additional factors mediate invasion and dissemination.

The IL-1 family are a large cohort of cytokines, receptors and adaptor proteins that play a broad and essential role in host immunology. They are categorised into three subgroups comprising IL-1/IL-33, IL-18 and IL-36, with signalling typically occurring when cytokines bind their receptor, recruit a co-receptor, and induce signalling through the Toll/interleukin-1 receptor domain resulting in mitogen activated protein kinase (MAPK) and nuclear factor kappa B (NF-κB) activation [10]. The IL-1 family have been implicated in the response to several fungal pathogens including *C. albicans*. IL-1α and IL-1β are induced during mucosal *C. albicans* infection and mediate acute (< 24 h) neutrophil recruitment [11] and drive the production of IL-17 [12]. IL-36α/β/γ are also induced during mucosal *C. albicans* infection and mice lacking IL-36R exhibit enhanced susceptibility to mucosal disease [12]. While the protective effect of IL-36 is not fully understood, IL-36R deficiency reduces IL-23 expression, a cytokine important for the proliferation and survival of IL-17 producing cells [12, 13]. IL-18 and IL-33 are induced by *C. albicans* infection [14, 15], though their roles in resolving mucosal disease remain unclear. In a mouse model of systemic candidiasis, IL-18 depletion increased disease susceptibility via an IFNγ-Th1 signalling axis [14]. Similarly, IL-33-deficient mice exhibited greater susceptibility to systemic infection, with IL-33 promoting IL-23 expression but downregulating IL-10 [15].

IL-1 family members are potent regulators of immunity, with insufficient activity frequently detrimental in a range of diseases but excessive activity being equally destructive [10, 16]. Here, we investigated how the combinatorial IL-1 family shapes the host immune response to mucosal *C. albicans* infection and explored the role of the IL- 1 family in mucosal-systemic dissemination. Using a murine model of IL-1 receptor accessory protein deficiency (IL-1RAcP^-/-^), we demonstrate that early protective immunity during oral *C. albicans* infection is critically dependent on IL-1 family signalling. Remarkably, inducing neutropenia in IL-1RAcP^-/-^ mice, but not wild type mice, results in dissemination of *C. albicans* to the liver and spleen (mimicking patient disease), and mortality. Our findings demonstrate that combined IL-1 family signalling is vital for host immunity against mucosal *C. albicans* infection and cooperates with neutrophils to prevent systemic dissemination. We identify the IL-1 family as a key host risk factor for fungal dissemination and suggest targeting IL-1 family signalling may prevent life threatening invasive infections in immunocompromised patients.

## Results

### The IL-1 family are rapidly induced at the mucosa and control fungal infection

To identify which IL-1 family members and immune recognition pathways were induced during *C. albicans* mucosal infection, we challenged TR146 human oral epithelial cells with *C. albicans* BWP17+CIp30 (wild-type; derivative of SC5314) and performed RNA-Sequencing. At 4 h post-infection, we observed the induction of IL-1α, IL-1β, IL-33 and IL-36γ but not toll-like receptor (TLR) and C-type lectin (CLR) receptor pathways (**fig. 1a and supplementary fig.1a**). Notably, IL-1 family members were some of the most significantly upregulated genes (**fig. 1b**). Given the rapid induction of IL-1 family signalling, and with IL-1α being a damage-associated cytokine, we hypothesised that IL- 1 family induction was dependent on the secretion of the damage-inducing peptide toxin candidalysin from *C. albicans* [5]. To test this, RNA-Sequencing was performed on TR146 cells infected with a *C. albicans* strain deficient in candidalysin (*ece1*Δ/Δ) and a re-integrant control (*ece1*Δ/Δ*+ECE1*), as well as synthetic candidalysin at weakly and strongly stimulating concentrations (3 µM and 15 µM; determined previously [5]). The induction of IL-1 family signalling was critically dependent on candidalysin (**fig. 1a and supplementary fig. 1b-d**), and the expression of IL-1 family members increased in a candidalysin concentration-dependent manner (**supplementary fig. 1e**).

**Figure 1:**
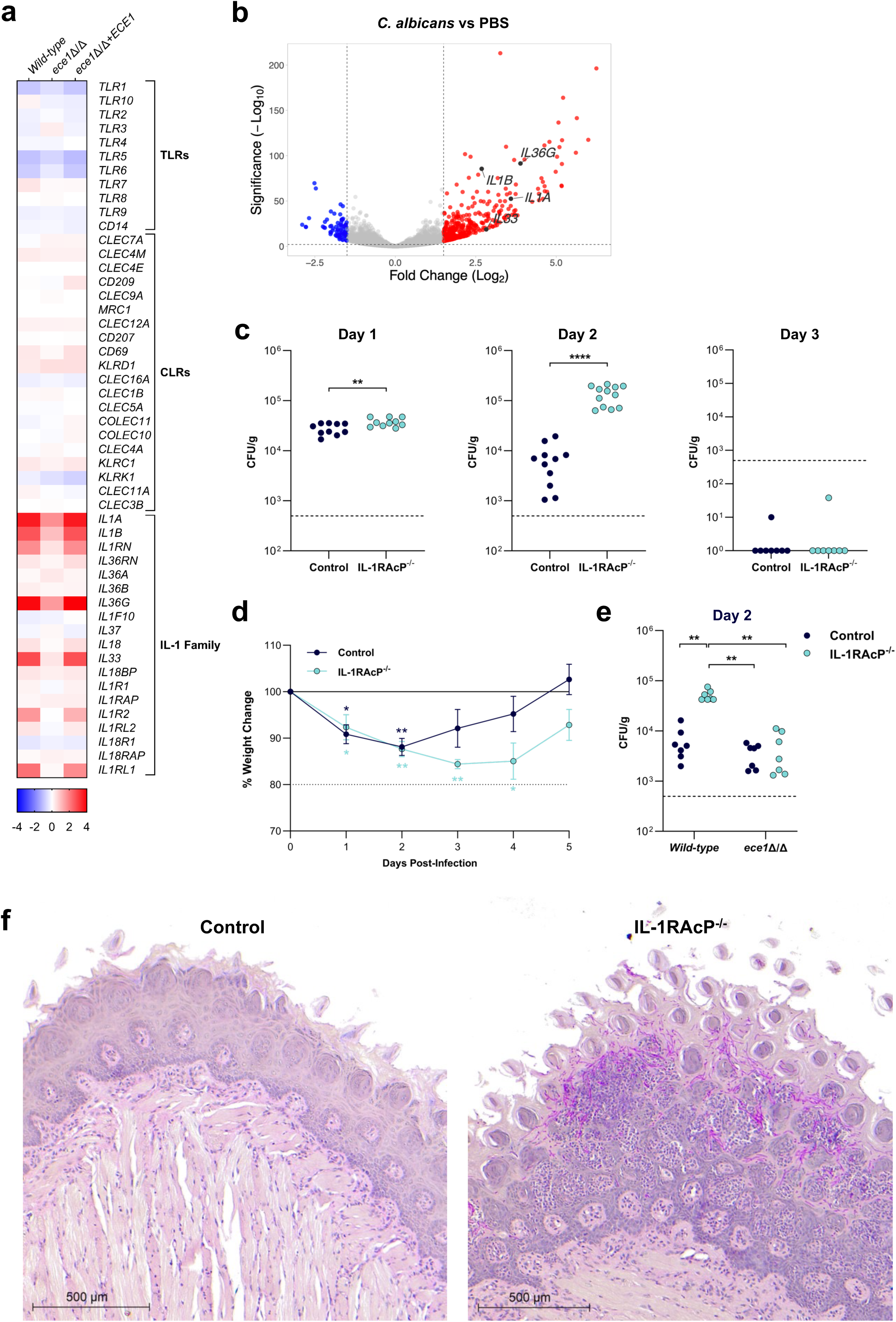
The IL-1 family are rapidly induced by candidalysin and restrict *C. albicans* infection and invasion. (**a-b**) TR146 cells were challenged with wild-type *C. albicans* (BWP17+CIp30), *ece1*Δ/Δ or *ece1*Δ/Δ+*ECE1* before bulk RNA sequencing was performed. (**a**) Selected constituents of TLR, CLR and IL-1 family signalling pathways were identified ([10, 56] and KEGG pathway reference hsa04620) and relative gene expression data displayed as a heat map. (**b**) Total bulk RNA gene expression data are displayed as a volcano plot according to their differential expression when compared to PBS. (**c-d**) Control (C57BL/6J) and IL-1RAcP^-/-^ mice were sublingually infected with wild type *C. albicans* (AHY940). (**c**) CFU was enumerated from tongue homogenate and analysed by Kruskal- Wallis test with Dunn’s post-test. Each symbol represents one mouse. Dotted line represents limit of detection. (**d**) Weight change of grouped control (dark blue line) and IL-1RAcP^-/-^ mice (light blue line) data from 3 independent experiments. Each timepoint was compared to the day 0 value by 2-way ANOVA with Bonferroni’s post-test. Dotted line represents experimental limit. (**e**) Control and IL-1RAcP^-/-^ mice were sublingually infected with *C. albicans* (AHY940) or *ece1*Δ/Δ. CFU was enumerated from tongue homogenate and analysed by 2-way ANOVA with Bonferroni’s post-test. Each symbol represents one mouse. Dotted line represents limit of detection. (**f**) Control and IL- 1RAcP^-/-^ mice were sublingually infected with *C. albicans* (AHY940) before tongue tissue was harvested on day 2 and Periodic acid-Schiff stained.

IL-1α, IL-1β, IL-33 and IL-36γ, the *C. albicans*-induced members of the IL-1 family, signal through a complex of their primary receptor (IL-1R, ST2 and IL-36R) and their common co-receptor, the IL-1 receptor accessory protein (IL-1RAcP). Therefore, we used an IL- 1RAcP^-/-^ murine model to investigate the impact of abolishing combinatorial IL-1 family signalling during *C. albicans* oral infection **(fig. 1a and supplementary fig. 1b-e**). We sublingually challenged IL-1RAcP^-/-^ and control (C57BL/6J) mice with *C. albicans* AHY940 (wild-type; derivative of SC5314) to induce oropharyngeal candidiasis (OPC), harvested tongue tissue, and quantified fungal burden (colony forming units (CFU)). IL- 1RAcP^-/-^ mice immediately (within 1 day) struggled to control the infection and displayed a remarkably acute susceptibility to a typically mild OPC challenge, as demonstrated by significantly higher fungal burdens at day 1 and most notably at day 2 (**fig. 1c**). Despite this, both control and IL-1RAcP^-/-^ mice cleared the infection by day 3. Weight change data corroborated the severity of the infection with IL-1RAcP^-/-^ mice experiencing significant weight loss for 4 days post infection (nearing the 80% experimental limit) before recovering (**fig. 1d**).

Given that the severity of OPC was dependent on IL-1 family signalling and IL-1 family induction was dependent on candidalysin, we next investigated whether candidalysin was driving virulence in the IL-1RAcP^-/-^ model. We challenged IL-1RAcP^-/-^ and control mice with *C. albicans* (AHY940) and *ece1*Δ/Δ to induce OPC and quantified CFU in tongue tissue. Notably, *ece1*Δ/Δ showed significantly reduced virulence in both control and IL-1RAcP^-/-^ mice (**fig. 1e and supplementary fig. 1f**), and there was no change in weight loss in either model (**supplementary fig. 1g**). This confirmed that candidalysin was driving *C. albicans* virulence in the IL-1RAcP^-/-^ model.

We next examined mucosal tissue to investigate the extent of *C. albicans* invasion. Histological sections of tongue tissue were taken from IL-1RAcP^-/-^ and control mice 2 days post-OPC and analysed with periodic acid-Schiff staining. While control mice had little detectable *C. albicans* and an intact mucosal barrier, tongue tissue from IL-1RAcP^-/-^ mice exhibited a heavily disrupted and inflamed mucosal barrier and contained large quantities of *C. albicans* hyphae, which invaded deep into the mucosa (**fig. 1f**). Our data demonstrate that combined IL-1 family signalling is critically important during early OPC and deficiencies in IL-1 signalling radically enhance susceptibility to *C. albicans* infection.

### Mucosal anti-microbial peptide and IL-17 expression require IL-1 family signalling

The production of AMPs and induction of innate type 17 immunity is critical for barrier immunity in acute OPC infections [6, 13, 17]. Thus, we investigated whether the high susceptibility of IL-1RAcP^-/-^ mice to OPC was associated with impairments in the Th17 axis. We induced OPC in IL-1RAcP^-/-^ and control mice, harvested mucosal tissue and quantified β-defensin 3, calprotectin (a heterodimer of S100a8 and S100a9), IL-17A/F, IL-22 and IL-23 gene expression. We found abolishment of β-defensin 3 expression and significantly reduced calprotectin expression at both day 1 and day 2 post infection in IL-1RAcP^-/-^ mice (**fig. 2a**). Given that β-defensin 3 is epithelial-derived, this suggests β-defensin 3 expression is entirely dependent on the IL-1 family-IL-17 axis during OPC. We also examined cathelicidin (*Camp*) but found no IL-1 family dependency (**supplementary fig. 2a**). IL-17A/F, IL-22 and IL-23 expression was largely dependent on IL-1 family signalling at day 1, but by day 2 (and with a fungal burden 10-times greater in the IL-1RAcP^-/-^ mice) the expression of IL-17A/F, IL-22 and IL-23 had fully recovered (**fig. 2b and supplementary fig. 2b**).

**Figure 2:**
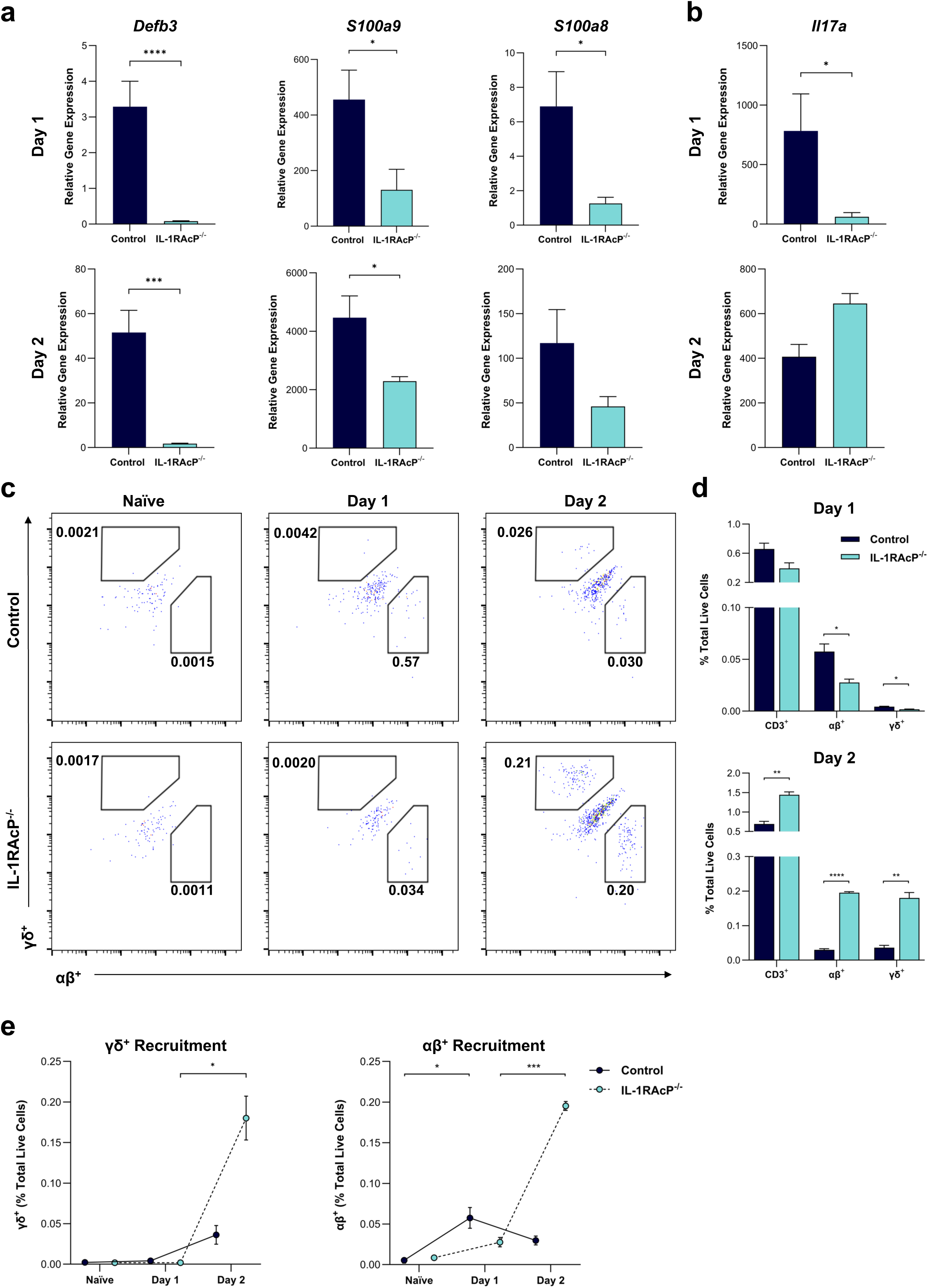
The IL-1 family initiate protective mucosal barrier immunity. (**a-e**) Control and IL-1RAcP^-/-^ mice were sublingually infected with *C. albicans* (AHY940). (**a-b**) RNA was extracted from tongue and gene expression determined by RT-qPCR. Results were quantified by ΔΔct analysis compared against a control naïve sample. Differences between control and IL-1RAcP^-/-^ mice were analysed by Kruskal-Wallis test with Dunn’s post-test. Graph displays cumulative data from at least 3 independent experiments (n=12 for each group). (**c-e**) Tongue tissue was homogenised to single cells and stained with a live/dead dye, anti-CD45, anti-CD3, anti-TCRαβ and anti-TCRγδ antibodies and/or their appropriate isotypes before being analysed by flow cytometry. (**c**) Flow plots representative of data from at least 3 independent experiments, gating shown as live (dead^+^ exclusion), single, CD45^+^, CD3^+^, and as percentage (%) of total live cells. (**d**) CD3^+^, αβ^+^ and γδ^+^ cells were quantified from at least 3 independent experiments (n=3 for each group) and differences between control and IL-1RAcP^-/-^ analysed by 2-way ANOVA with Bonferroni’s post-test. (**e**) αβ^+^ and γδ^+^ cells (as % total live cells) from at least 3 independent experiments (n=3 for each group) were tracked from naïve to day 2 of infection. Differences within each group between each timepoint were analysed by 2-way ANOVA with Bonferroni’s post-test.

The production of IL-17 during acute OPC is predominantly mediated by innate acting ‘natural’ TCRαβ^+^ and TCRγδ^+^ cells [13, 18]. Therefore, we induced OPC in IL-1RAcP^-/-^ and control mice and assessed the infiltration and expansion of TCRαβ^+^ and TCRγδ^+^ cells. While naïve (uninfected) mice exhibited no difference in cellular infiltration (**supplementary fig. 2c,d**), the acute infiltration (day 1) of TCRαβ^+^ and TCRγδ^+^ cells was IL-1 family dependent (**fig. 2c,d**). However, by day 2, cellular infiltration had fully recovered at levels far greater than control mice (**fig. 2e**), indicating that cellular recruitment once infection was established was IL-1 family independent. These data suggest that early protective barrier immune responses in OPC require combined IL-1 family signalling, and their absence permits *C. albicans* overgrowth and increased virulence.

### Mucosal neutrophil infiltration is delayed without IL-1 family signals

Neutrophil recruitment is a defining feature of OPC and we observed high quantities of inflammatory infiltrate in IL-1RAcP^⁻/⁻^ mice 2 days after OPC (**fig. 1f**). To investigate neutrophil recruitment, we induced OPC in IL-1RAcP^-/-^ and control mice, harvested tongue tissue and stained histological sections with hematoxylin and eosin. Interestingly, extensive neutrophil infiltration was observed only in IL-1RAcP^-/-^ at day 2 (**fig. 3a**). Next, to explore how the IL-1 family influences the recruitment of neutrophils to the mucosa, we quantified the expression of *Cxcl1* and *Csf3* (key neutrophil recruiting chemokines). The rapid induction (day 1) of both *Cxcl1* and *Csf3* was highly dependent on the IL-1 family; however, by day 2 both genes were potently induced in the IL-1RAcP^-/-^ model (**fig. 3b and supplementary fig. 3a**). Strikingly, using flow cytometry, neutrophil recruitment in IL-1RAcP^-/-^ mice was 10-fold lower on day 1. However, by day 2, recruitment surged dramatically, with neutrophil numbers increasing 100-fold compared to day 1 and exceeding those in control mice by 7-fold (**fig. 3c-e and supplementary fig. 3b,c**). To further explore this neutrophil infiltration, we examined emergency granulopoiesis by assessing changes in bone marrow composition (**supplementary fig. 3d,e**). Although a modest release of mature neutrophils was observed in control mice, we identified a large but delayed release of neutrophils in IL- 1RAcP^-/-^ mice at day 2. Our data demonstrate that the early recruitment of neutrophils during OPC is dependent on IL-1 family signalling, but once OPC becomes severe IL-1 family independent mechanisms provide redundancy.

**Figure 3:**
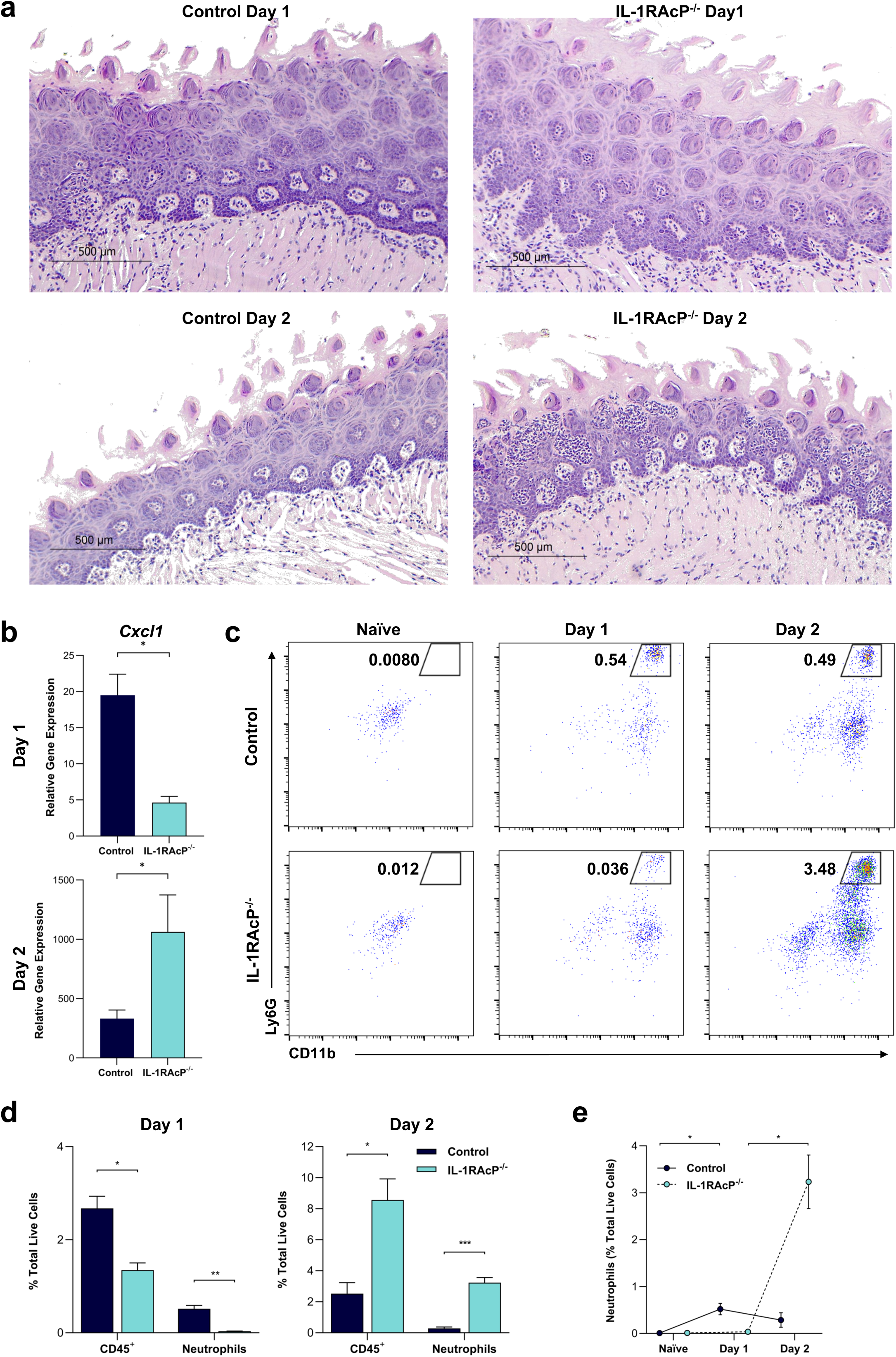
Rapid mucosal neutrophil recruitment is dependent on IL-1 family signalling. (**a-e**) Control and IL-1RAcP^-/-^ mice were sublingually infected with *C. albicans* (AHY940). (**a**) Tongue tissue was harvested and stained with hematoxylin and eosin. (**b**) RNA was extracted from tongue and gene expression determined by RT-qPCR. Results were quantified by ΔΔct analysis compared against a control naïve sample. Graph displays cumulative data from at least 3 independent experiments (n=12 for each group). Differences between control and IL-1RAcP^-/-^ were analysed by Kruskal-Wallis test with Dunn’s post-test. (**c**) Tongue tissue was homogenised to single cells and stained with a live/dead dye, anti-CD45, anti-Ly6G, and anti-CD11b antibodies and/or their appropriate isotypes before being analysed by flow cytometry. Flow plots representative of data from at least 3 independent experiments, gating shown as live (dead^+^ exclusion), single, CD45^+^, and as percentage (%) of total live cells. (**d**) CD45^+^ and CD45^+^ Ly6G^+^ CD11b^+^ cells were quantified from at least 3 independent experiments (n=3 for each group). Differences between control and IL-1RAcP^-/-^ were analysed by 2-way ANOVA with Bonferroni’s post-test. (**e**) CD45^+^ and CD45^+^ Ly6G^+^ CD11b^+^ cells (as % total live cells) from at least 3 independent experiments (n=3 for each group) were tracked from naïve to day 2 of infection. Differences within each group at each timepoint were analysed by 2-way ANOVA with Bonferroni’s post-test.

### Mucosal-systemic dissemination is governed by the IL-1 family and neutrophils

Our data demonstrate that IL-1 family signalling is critical for the initial control of OPC infection (day 1), but that even without IL-1 family signalling neutrophils are eventually recruited (day 2) and infection is resolved. We therefore hypothesised that IL-1 family signalling and neutrophils cooperate to provide protection against OPC as the infection progresses. To test this, we depleted neutrophils (**supplementary fig. 4a,b**) in both IL- 1RAcP^-/-^ and control mice using anti-Ly6G antibodies and induced OPC. Strikingly, we observed severe mucosal infection in neutropenic IL-1RAcP^-/-^ mice which succumbed to the infection by day 4, confirming that neutrophils provide potent redundancy in the absence of IL-1 family signalling (**fig. 4a and supplementary fig. 4c**). In contrast, neutropenic (wild type) control mice largely cleared the infection with minimal weight loss (**fig. 4b and supplementary fig. 4c**). Also, while *C. albicans* could be recovered from the stomach, colon and cecum (via swallowing) in both neutropenic control and neutropenic IL-1RAcP^-/-^ mice at similar levels (**supplementary fig. 4d**), the fungus could only be recovered from internal organs of neutropenic IL-1RAcP^-/-^ mice. *C. albicans* initially targeted the liver (day 3) but also spleen, kidney and brain (day 4) (**fig. 4c,d**). Importantly, the liver was the predominant target of disseminated *C. albicans*, mimicking the primary organ *C. albicans* targets in severely immunocompromised patients [4, 19, 20]. Our data demonstrate that IL-1 family signalling and neutrophils cooperate to limit fungal translocation across mucosal barriers and prevent subsequent life-threatening systemic infections.

**Figure 4:**
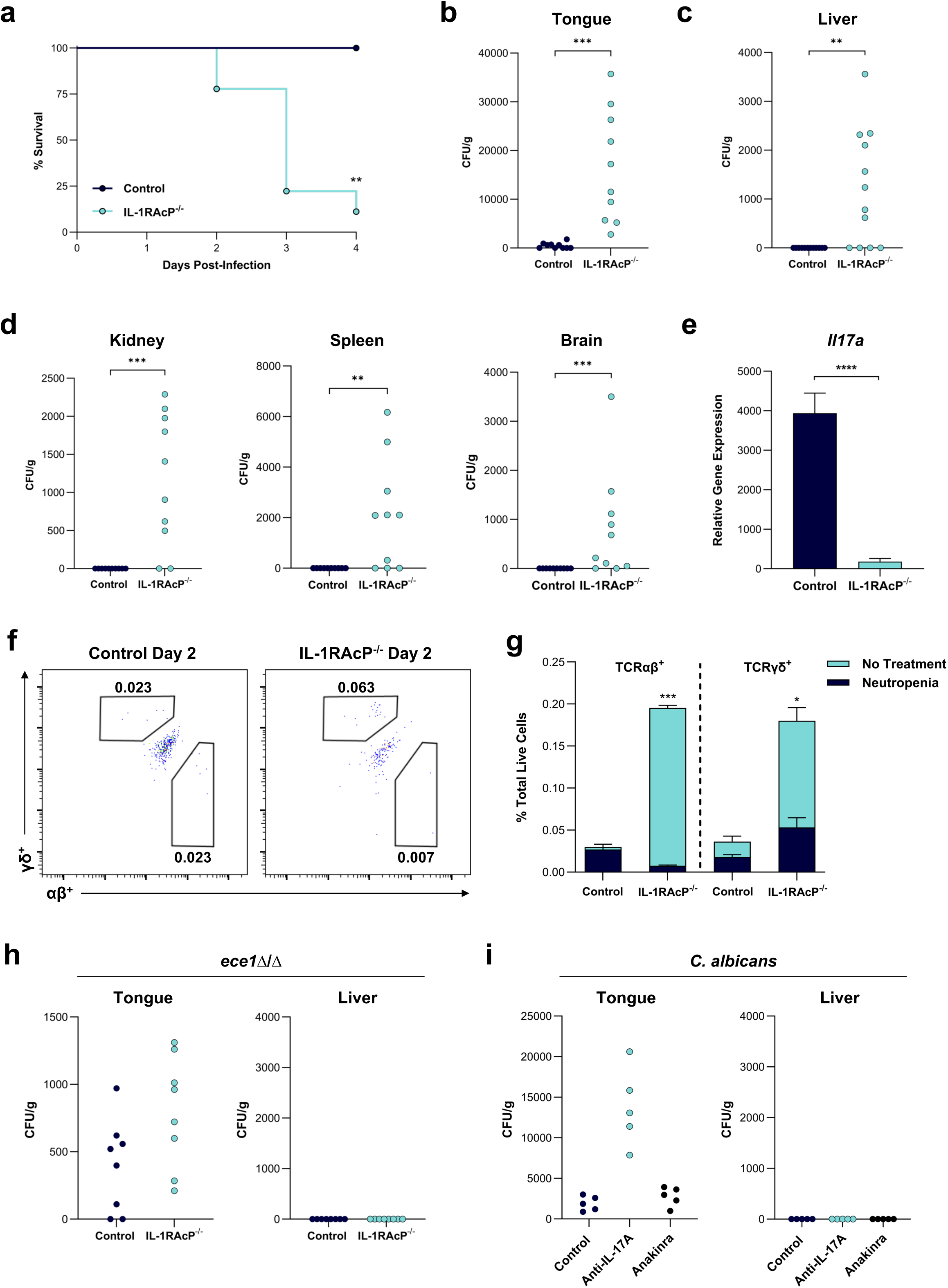
The IL-1 family and neutrophils restrict invasion and restrain *C. albicans* within the mucosa. (**a-g**) Neutrophils were depleted in control and IL-1RAcP^-/-^ mice by intraperitoneal administration of anti-Ly6G antibody starting 1 day prior to sublingual administration of *C. albicans* (AHY940). (**a**) Mice were weighed and scored daily and any mice exceeding experimental severity limits were culled. Graph displays cumulative data from at least 3 independent experiments (n=12 for each group). Survival differences between control and IL-1RAcP^-/-^ were analysed by log-rank test. (**b**) CFU was enumerated from tongue homogenate at day 4 (**c**) CFU was enumerated from liver homogenate at day 3. (**d**) CFU was enumerated from kidney, spleen and brain homogenate at day 4. (**b-d**) Each symbol represents one mouse. CFU differences were analysed by Kruskal-Wallis test with Dunn’s post-test. (**e**) RNA was extracted from tongue on day 2 and gene expression determined by RT-qPCR. Results were quantified by ΔΔct analysis compared against a naïve sample. Graph displays cumulative data from at least 3 independent experiments (n=6 for each group). Differences between control and IL-1RAcP^-/-^ were analysed by Kruskal-Wallis test with Dunn’s post-test. (**f**) Tongue tissue was homogenised to single cells and stained with a live/dead dye, anti-CD45, anti-CD3, anti-TCRαβ and anti-TCRγδ antibodies and/or their appropriate isotypes before being analysed by flow cytometry. Flow plots representative of data from at least 3 independent experiment and gating shown as live (dead^+^ exclusion), single, CD45^+^, CD3^+^, and as percentage (%) of total live cells. (**g**) CD3^+^, αβ^+^ and γδ^+^ cells were quantified from at least 3 independent experiments (n=3 for each group) and differences between neutropenic and non- neutropenic for control and IL-1RAcP^-/-^ mice were analysed by 2-way ANOVA with Bonferroni’s post-test. (**h**) Neutrophils were depleted in control and IL-1RAcP^-/-^ mice by intraperitoneal administration of anti-Ly6G antibody starting 1 day prior to the sublingual administration of *C. albicans ece1*Δ/Δ. CFU was enumerated from tongue and liver homogenate at day 4. Each symbol represents one mouse. CFU differences were analysed by Kruskal-Wallis test with Dunn’s post-test. (**i**) Neutrophils were depleted in control mice by intraperitoneal administration of anti-Ly6G antibody starting 1 day prior to sublingual administration of *C. albicans* (AHY940). Anti-IL-17A antibody or Anakinra was administered by intraperitoneal administration starting 1 day prior to sublingual administration of *C. albicans* (AHY940). CFU was enumerated from tongue and liver homogenate at day 4. Each symbol represents one mouse. CFU differences were analysed by Kruskal-Wallis test with Dunn’s post-test.

Since sufficient protection against *C. albicans* dissemination was provided by either the IL-1 family or neutrophils, we next investigated whether neutrophils were responsible for the delayed recruitment of TCRαβ^+^ and TCRγδ^+^ cells in IL-1RAcP^-/-^ mice at day 2 post infection. Interestingly, we found that depletion of neutrophils blocked IL-17A expression and TCRαβ^+^ and TCRγδ^+^ infiltration (**fig. 4e-g**). While the expression of the IL- 17-associated genes IL-22 and IL-23 was also reduced, the expression of neutrophil attracting chemokines was significantly increased despite the absence of neutrophils (**supplementary fig. 4e**).

Candidalysin is vital for *C. albicans* infection, and critically, for the induction of mucosal immunity through IL-1 family signalling [5–7, 21]. To determine whether candidalysin was required for fungal dissemination, we challenged neutropenic IL-1RAcP^-/-^ and control mice with wild-type *C. albicans* (AHY940) and *ece1*Δ/Δ. Despite both fungal strains being equally capable of colonising mucosal tissues, *ece1*Δ/Δ was unable to translocate across mucosal barriers and was absent from internal organs including the liver, demonstrating that candidalysin is essential for fungal dissemination (**fig. 4h and supplementary fig. 5 a,b**).

Finally, we questioned whether IL-1 family members together controlled dissemination or whether IL-1R deficiency or IL-17 deficiency alone permitted dissemination. To test this, we challenged neutropenic control mice with anakinra (IL-1R antagonist) or anti-IL- 17A antibodies and infected with wild-type *C. albicans* (AHY940). Anakinra had little effect, with mice displaying low oral fungal burden, no fungal dissemination, and weight loss recovering from day 3. Similarly, while anti-IL-17A antibodies rendered neutropenic mice more susceptible to persistent infection and greater weight loss, no fungal dissemination was observed (**fig. 4i and supplementary fig. 5c**). Together, our data demonstrate that the combined IL-1 family cooperates with neutrophils to prevent mucosal dissemination of *C. albicans*. Therefore, this study identifies the IL-1 family as a key host risk factor for life threatening disseminated fungal disease.

## Discussion

Mechanisms that control microbial overgrowth to prevent mucosal disease and limit dissemination within the host are poorly understood. Fungal infections are a serious threat to global health, particularly as the number of immunocompromised patients is increasing [22]. With limited therapeutic options and antifungal resistance on the rise, there is an urgent need to identify host factors capable of controlling mucosal infection and preventing dissemination. Establishing host risk factors that enhance susceptibility to invasive fungal disease may permit a personalised medicine strategy in high-risk patients and the potential therapeutic reinforcement of these factors. Our results identify a central role for the IL-1 family in mediating rapid and protective immunity against mucosal *C. albicans* challenge. Crucially, the IL-1 family can prevent mucosal *C. albicans* infection from disseminating in the absence of neutrophils. Similarly, in the absence of the IL-1 family, a delayed neutrophil response eventually resolves mucosal *C. albicans* infection and prevents dissemination. Critically, absence of IL-1 family signalling coupled with neutropenia permits the dissemination of *C. albicans* from the mucosa, first to the liver and then into multiple organs, mimicking disease experienced by severely immunocompromised patients. Our data identifies the IL-1 family as a key host risk factor for life threatening disseminated disease.

The activation of host mucosal immunity by *C. albicans* through candidalysin-induced cellular damage is well described [5, 8, 12, 23]. In agreement, we confirmed that the induction of IL-1α, IL-1β, IL-33 and IL-36γ was candidalysin dependent. Although clear and important anti-fungal roles for CLRs and TLRs have been described [24, 25], we observed no gene induction of either receptor family, corroborating their lack of involvement in driving rapid oral mucosal immunity during OPC [6, 26]. Modelling IL-1 family deficiency *in vivo* using IL-1RAcP^-/-^ mice revealed a high susceptibility to OPC infection with deep and widespread hyphal invasion, which typically requires extensive immune suppression to reach such severity [27]. OPC infection was driven by candidalysin as *C. albicans* deficient in candidalysin (*ece1*Δ/Δ) was unable to establish OPC even in IL-1RAcP^-/-^ mice.

The barrier immune response against mucosal pathogens, including *C. albicans*, relies heavily on AMP production [28]. In patients, a defective AMP compartment leads to reduced candidacidal capacity and elevated *C. albicans* colonisation [29]. While candidalysin promotes the induction of key AMPs including β-defensins and calprotectin [7], our results show the host drives this response through IL-1 family signalling. Interestingly we found cathelicidin expression was independent of IL-1 family signalling, which agrees with a previous study showing that cathelicidin expression does not require candidalysin [7]. IL-17 plays a pivotal role in AMP production by mediating the activation of IκBζ which drives β-defensin 3 expression [17]. Critically, candidalysin was again identified as a key driver of this signalling pathway. Thus, our data suggests that the IL-1 family sits upstream of IL-17 in this pathway. AMPs are critical for limiting the initial growth of pathogenic *C. albicans* [30], and we hypothesise abolishing the IL-1 family dependent AMP response permits *C. albicans* overgrowth.

In addition to AMPs, the expression of IL-17 is central to barrier protection. Defects in the IL-17 pathway enhance both mucosal and systemic *C. albicans* susceptibility in mice and humans [13, 31, 32]. IL-17 has been linked to neutrophil recruitment, with studies suggesting IL-17 assists neutrophil infiltration but recruitment is not IL-17- dependent [33, 34]. IL-17 is predominantly produced in response to mucosal *C. albicans* infection by γδ T cells and natural Th17 cells (characterised by their expression of TCRαβ) and requires IL-23 [35]. Our results reveal that IL-1 family signalling drives both the rapid recruitment of γδ T cells and natural Th17 cells, and the early production of IL-17. Previous studies using IL-1R and IL-36R deficient models indicate both pathways independently but only partially control IL-17 production [6, 12]. It was thought that natural Th17 cells alone expand during acute OPC infection [6]; however, we identified increased γδ T cells in the IL-1RAcP^-/-^ model. Given that we also observe a delayed increase in natural Th17 cells, we hypothesise that the severity of the mucosal infection initiates the IL-1 family-independent restoration of IL-17 production through both γδ T and natural Th17 cells. While γδ T cells are known to infiltrate into other tissues during disease [36], even during peritoneal candidiasis [37], their role in mucosal *C. albicans* infection requires further investigation.

Neutrophils comprise a key arm of the mucosal immune response and when recruited provide potent mucosal protection [38]. The recruitment of neutrophils during OPC requires IL-1R signalling [11]. In agreement, we found both chemokine expression and neutrophil recruitment were dependent on the IL-1 family. However, the eventual restoration of chemokine expression and neutrophil infiltration was IL-1 family independent. Interestingly, we also identified an influx of immature neutrophils, suggesting not only that emergency granulopoiesis can be initiated independently of the IL-1 family, but that the IL-1RAcP^-/-^ model required substantial neutrophil infiltration to combat severe OPC infection. Taken together, our data suggest the IL-1 family plays a critical role in the mucosa controlling the release of AMPs, early IL-17 expression, and rapid neutrophil recruitment. IL-1RAcP^-/-^ mice were highly susceptible to OPC but infection was eventually resolved through the IL-1 family independent recovery of IL-17 expression and extensive infiltration of neutrophils.

Neutrophils are potent anti-fungal immune cells, particularly due to their ability to kill physically large pathogens including *C. albicans* hyphae [39], and neutropenia is the most widely accepted patient risk factor for invasive fungal disease [40]. Given this, we hypothesised neutrophils were resolving the severe infection in the IL-1RAcP^-/-^ model. In agreement, inducing neutropenia in IL-1RAcP^-/-^ mice resulted in a severe OPC infection, and mortality within four days. Remarkably, we found that *C. albicans* translocated across mucosal barriers to invade the liver (day 3) and subsequently the spleen, kidney and brain (day 4), mimicking the dissemination of *C. albicans* to multiple organs in human disease [4, 19]. This is particularly noteworthy as modelling dissemination *in vivo* has proven challenging. Current systemic *C. albicans* infection models, which are initiated via intravenous injection and lead to kidney infection, fail to capture the critical mucosal events governing dissemination in humans.

Strikingly, we found that inducing neutropenia in IL-1RAcP^-/-^ mice also eliminated their ability to recover IL-17 expression or to recruit γδ T cells and natural Th17 cells. Neutrophils can produce IL-17 [41, 42] and our results suggest that infiltrating neutrophils may crosstalk with the IL-17 axis. While this interaction has been previously described [43], it requires further investigation during OPC. Also, although IL-17 deficiency can result in chronic mucosal infection [44] and enhances susceptibility to systemic infection [31, 45], we corroborate that blocking IL-17 in neutropenic wild type mice does not result in dissemination [13]. This suggests that combinatorial IL-1 family signalling, rather than IL-17 alone, is the key immune factor restricting fungal dissemination across the mucosa.

In summary, this study reveals that deficiencies in IL-1 family signalling permit uncontrolled mucosal *C. albicans* infection which, when coupled with neutropenia, results in mucosal-systemic dissemination and mortality. While antibiotics, chemotherapeutic agents and dextran sodium sulphate are typically used to promote the dissemination of microbes [46], we identify the IL-1 family as a host factor critical for the control of mucosal-systemic dissemination of an invasive fungal pathogen. Notably, patients with polymorphisms leading to deficient IL-1 family signalling are more susceptible to invasive candidiasis [47] and disseminated fungal infection [48]. Previous studies have retrospectively stratified patient cohorts according to their immune function and their risk of acquiring fungal disease, but this has not been undertaken for the IL-1 family [49–51]. Our findings suggest that combinatorial IL-1 family function plays a crucial role in dissemination risk, offering potential for a personalised therapeutic approach. Consequently, therapeutically enhancing IL-1 family function to augment mucosal immunity and reduce dissemination could have significant clinical implications.

## Methods

### *Candida albicans* strains and culture

BWP17+CIp30 (wild-type), *ece1*Δ/Δ (*ECE1*-deficient), and *ece1*Δ/Δ+*ECE1* (*ECE1* revertant), (all previously described [15]) were used for TR146 infection and RNA sequencing analysis. All subsequent data from the study used AHY940 (SC5314 a/α *leu2*Δ/*LEU2*), or *ece1*Δ/Δ (*leu*2Δ/*LEU2 ece1*ΔΔ), an *ECE1* ORF deletion mutant constructed using CRISPR-Cas9 in parental strain AHY940 [52]. All *C. albicans* strains were plated on YPD (yeast extract peptone dextrose) agar, cultured for 16–20 h in YPD broth, washed three times with PBS and resuspended at the required concentration in PBS.

### TR146 cell culture, treatment and RNA sequencing

TR146 human buccal epithelial squamous cell carcinoma cells acquired from the European Collection of Authenticated Cell Cultures were cultured in Dulbecco’s modified Eagle medium/Nutrient Mixture F-12 (DMEM/F12) (Gibco, ThermoFisher Scientific) supplemented with 10% (v/v) foetal bovine serum (FBS; Gibco, ThermoFisher Scientific) and 1% penicillin-streptomycin (Gibco, ThermoFisher Scientific) at 37°C 5% CO2. Prior to treatment, confluent TR146 cells were serum-starved overnight in DMEM/F12 medium.

For RNA-Seq, TR146 cells were infected with *C. albicans* strains or PBS for 4 h at an MOI (multiplicity of infection) of 10 (10 fungal cells to 1 epithelial cell), or 3 or 15 µM candidalysin in deionised water or a deionised water control for 6 h. Total RNA from TR146 monolayers was isolated using the Nucleospin II kit (Macherey-Nagel, ThermoFisher Scientific). RNA quality control was performed with an Agilent RNA 600 Nano Kit using the Bioanalyzer. Libraries were prepared using a TruSeq RNA Sample Preparation kit V2 (Illumina). Paired-end raw sequencing data (FASTQ files) were aligned to human reference (GRCh38) from ensemble release 92 using “Kallisto” software [53] to quantify Transcripts Per Million (TPM) and count values of the transcripts. The gene-level count and TPM values were calculated from the transcript-level abundance and counts using R package “tximport” [54]. Differential expression analyses were performed based on gene-level counts (having Entrez ID and the median TPM>1 across all samples) using R package “DESeq2” and Wald test p value (FDR<0.01). R package “org.Hs.eg.db” was used for ID mapping and Hierarchical clustering with Ward.D2 and Euclidean distance were used to cluster the samples based on the similarity matrix. Additionally, principal component analysis (PCA) that visualised using R package “ggplot2” was performed based on the log transformed TPMs. Volcano plots were generated with VolcaNoseR ([55]).

### *In vivo* murine studies

Mice were maintained and used according to institutional and UK Home Office guidelines. This study was performed in strict accordance with the Project License (PP6711863). The animal care and use protocol adhered to the Animals (Scientific Procedures) Act 1986. B6;129S1-Il1raptm1Roml/J were purchased from Jax (RRID:IMSR_JAX:003284) and backcrossed for 10 generations to C57BL/6J (RRID:IMSR_JAX:000664). Throughout the study IL1RAcP^-/-^ represent the backcrossed B6;129S1-Il1raptm1Roml/J, and control mice represent C57BL/6J. Male and female 8-12 week old mice were used in experimentation.

### Murine oropharyngeal candidiasis model

OPC was induced as previously described [27]. Briefly, mice were anaesthetised for 75 min with an intra-peritoneal injection of 75 mg/kg ketamine and 1 mg/kg domitor. A swab soaked in a 10^7^ CFU/ml *C. albicans* yeast in sterile PBS was placed sub-lingually for 75 min. After 1, 2, 3 or 4 days, mice were sacrificed, the tongue excised and divided longitudinally in half. One half was weighed, homogenised using a Gentle MACS (Miltenyi Biotec) and serial dilutions were plated on YPD agar containing 50 μg/ml chloramphenicol. The plates were cultured at 37°C for 24 h and CFU was calculated per g organ. The other half was processed for RNA extraction, flow cytometry or histochemistry.

### Histology

Mouse tongue tissue was collected, formalin fixed, dehydrated and paraffin embedded before 5 micrometer sections were cut and mounted on slides. Slides then underwent Periodic Acid Shift staining or hematoxylin and eosin staining following standard protocols. Briefly for PAS staining, sections were treated with 1% periodic acid for 10 minutes followed by Schiff reagent for 10 minutes before counterstaining with Harris Hamotoxylin. Briefly for HCE staining, sections were treated with Harris Hamotoxylin before washing with water and counterstaining with eosin.

### Gene expression analysis

Half tongue was collected into RNALater (Thermo Fisher Scientific) and homogenised using a Gentle MACS (Miltenyi Biotec). Total RNA was harvested using an RNeasy Kit QIAGEN). cDNA was generated using the QuantiTect Rev. Transcription Kit (QIAGEN). qRT-PCR was performed using a QuantiNova SYBR Green PCR kit (QIAGEN) with predesigned primer pairs (**Supplementary Table 1**) purchased from Merck. qRT-PCR was run on a QuantStudio 7 (Thermo Fisher Scientific). Gene expression was calculated using ΔΔCt where a control (typically naïve) sample’s experimental value was set at 1.

### Tongue Digestion and Flow cytometry

Half tongue was collected on ice and homogenised using a Gentle MACS (Miltenyi Biotec) with a tumour dissociation kit (Miltenyi Biotec). Single cell homogenate was passed through a 70 µm filter and washed with pre-cooled RPMI 1640 without phenol red (Gibco, ThermoFisher) before being counted and resuspended in pre-cooled PBS. Cells were live/dead stained with Zombie Green Fixable Viability Kit (Biolegend) for 30 min at 4°C and then washed in pre-cooled FACS Buffer (PBS 3% FBS 2 mM EDTA). Cells were blocked with TruStain FcX (anti-mouse CD16/32) (Biolegend) for 15 min at 4°C before being stained with the required antibodies and/or isotypes (**Supplementary Table 1**) for 30 min at 4°C. Cells were washed three times with pre-cooled FACS buffer before being run on a BD FACSCanto II (BD Biosciences). Results were analysed using FlowJo software (BD Biosciences).

### Induction of neutropenia, anti-IL-17A treatment, and anakinra treatment

Neutropenia was induced through the intraperitoneal injection of murinised anti-Ly6G antibody (Absolute Antibodies, Absolute Biotech). On day −1 a loading dose of 300 µg/mouse was administered. Maintenance doses of 150 µg/mouse were administered on day 1 and day 3. Isotype control was administered at the same doses and time points. Neutropenia was confirmed in tongue tissue using flow cytometry. Neutropenia was also confirmed in blood extracted by cardiac puncture. Briefly, extracted blood was collected into PBS containing 4 mM EDTA and red blood cells lysed using RBC lysis buffer (Biolegend). Cells were washed with PBS, blocked with TruStain FcX (anti-mouse CD16/32) for 15 min at 4°C and stained with the required antibodies and/or isotypes (**Supplementary Table 1**) for 30 min at 4°C. Cells were washed three times with pre- cooled FACS buffer before being run on a BD FACSCanto II (BD Biosciences). Results were analysed using FlowJo software (BD Biosciences).

IL-17A was blocked through intraperitoneal administration of InVivoMAb anti-IL-17A antibody (Bio X Cell). A dose of 200 µg/mouse was administered on day −1, 1 and 3. Isotype control was administered at the same doses and time points. IL-1R was blocked through intraperitoneal administration of anakinra (KINERET, Sobi). A dose of 200 µg/mouse was administered on day −1, 1 and 3.

### Organ harvest

Whole brain, liver, kidney, spleen, stomach and cecum were harvested and weighed. A 2 cm section of colon was harvested and weighed. All tissue was subsequently homogenised using a Gentle MACS (Miltenyi Biotec) and serial dilutions were plated on YPD agar containing 50 μg/ml chloramphenicol. The plates were cultured at 37°C for 24 h and CFU was calculated per g organ.

### Statistical analysis

Data are presented as mean or median +/- S.E.M. and are representative or cumulative data from the indicated number of independent experiments. Data was tested for normality and if data followed a Gaussian distribution a one-way ANOVA followed by Bonferroni’s post-test or 2-way ANOVA followed by Bonferroni’s post-test was used when multiple groups were analysed. A Kruskal-Wallis test with Dunn’s post-test was used when non parametric data were analysed. A Gaussian distribution was assumed for experiments with small sample numbers. Data containing zero values was transformed by Y = sqrt(Y+0.5). Statistical significance was set at *p<0.05, **p<0.005, ***p<0.001, ****p<0.0001.

## Figures

**Supplementary figure 1**

(**a-d**) TR146 cells were challenged with wild-type *C. albicans* (BWP17+CIp30) or *ece1*Δ/Δ before bulk RNA sequencing was performed. (**a-b**) Log_2_FC values for IL-1 family members were identified after challenge with either *C. albicans* (BWP17+CIp30) or *ece1*Δ/Δ and their significance analysed. (**c-d**) Total bulk RNA gene expression results for *ece1*Δ/Δ or *ece1*Δ/Δ+*ECE1* displayed as a volcano plot according to their differential expression when compared to PBS. (**e**) TR146 cells were challenged with 3 µM or 15 µM candidalysin before bulk RNA sequencing was performed. Log_2_FC values for IL-1 family members were calculated and their significance analysed. (**f-g**) Control and IL-1RAcP^-/-^ mice were sublingually infected with *C. albicans* (AHY940) or *ece1*Δ/Δ. (**f**) CFU was enumerated from tongue homogenate and analysed by 2-way ANOVA with Bonferroni’s post-test. Each symbol represents one mouse. (**g**) Weight change of grouped control (dark blue line) and IL-1RAcP^-/-^ mice (light blue line) data from 2 independent experiments (n=7 for each group). Each timepoint was compared to the day 0 value by 2-way ANOVA with Bonferroni’s post-test. Black dotted line represents experimental limit.

**Supplementary figure 2**

(**a-c**) Control and IL-1RAcP^-/-^ mice were sublingually infected with *C. albicans* (AHY940). (**a-b**) RNA was extracted from tongue and gene expression determined by RT-qPCR. Results were quantified by ΔΔct analysis compared against a naïve sample. Differences between control and IL-1RAcP^-/-^ were analysed by Kruskal-Wallis test with Dunn’s post-test. Graph displays cumulative data from at least 3 independent experiments (n=12 for each group). (**c-d**) Tongue tissue was homogenised to single cells and stained with a live/dead dye, anti-CD45, anti-CD3, anti-TCRαβ and anti-TCRγδ antibodies and/or their appropriate isotypes before being analysed by flow cytometry. (**c**) Gating strategy for isolating CD3^+^, αβ^+^ and γδ^+^ cells completed on day 2 post-OPC IL-1RAcP^-/-^ sample. (**d**) CD3^+^, αβ^+^ and γδ^+^ cells were quantified from at least 3 independent experiments (n=3 for each group) and differences between control and IL-1RAcP^-/-^ analysed by 2-way ANOVA with Bonferroni’s post-test.

**Supplementary figure 3**

**(a)** Control and IL-1RAcP^-/-^ mice were sublingually infected with *C. albicans* (AHY940). RNA was extracted from tongue and gene expression determined by RT-qPCR. Results were quantified by ΔΔct analysis compared against a naïve sample. Graph displays cumulative data from at least 3 independent experiments (n=12 for each group). Differences between control and IL-1RAcP^-/-^ were analysed by Kruskal-Wallis test with Dunn’s post-test. (**b**) CD45^+^ and CD45^+^ Ly6G^+^ CD11b^+^ cells were quantified from at least 3 independent experiments (n=3 for each group). Differences between control and IL- 1RAcP^-/-^ were analysed by 2-way ANOVA with Bonferroni’s post-test. (**c-f**) Control and IL- 1RAcP^-/-^ mice were sublingually infected with *C. albicans* (AHY940). (**c**) Gating strategy for isolating CD45^+^ Ly6G^+^ CD11b^+^ cells completed on day 2 post-OPC control sample. (**d-e**) Bone marrow was harvested from control and IL-1RAcP^-/-^ femurs, single cells isolated and stained with anti-CD45, anti-Ly6G, anti-Ly6C, and anti-CD11b antibodies and/or their appropriate isotypes before being analysed by flow cytometry. CD45^+^ CD11b^+^ Ly6G^HI^ and Ly6C^INT^ cells were identified as mature neutrophils. (**d**) Flow plots representative of data from at least 3 independent experiments, gating shown as single, CD45^+^, CD11b^+^ and as percentage (%) of total CD45^+^ CD11b^+^ cells. (**e**) CD45^+^ CD11b^+^ Ly6G^HI^ and Ly6C^INT^ cells were quantified from at least 3 independent experiments (n=3 for each group). Differences between control and IL-1RAcP^-/-^ at each timepoint were analysed by 2-way ANOVA with Bonferroni’s post-test. (**f**) Gating strategy for isolating CD45^+^ CD11b^+^ Ly6G^HI^ and Ly6C^INT^ cells completed on day 2 post-OPC control sample.

**Supplementary figure 4**

**(a)** Blood was drawn from control mice 1 day after intraperitoneal administration of anti- Ly6G antibody or isotype control and stained with anti-CD45, anti-Ly6G, and anti- CD11b antibodies before being analysed by flow cytometry. Flow plots representative of data from 2 independent experiments, gating shown as single CD45^+^ and as percentage (%) of total CD45^+^ cells. (**b-e**) Neutrophils were depleted in control and IL-1RAcP^-/-^ mice by intraperitoneal administration of anti-Ly6G antibody starting 1 day prior to sublingual administration of *C. albicans* (AHY940). (**b**) After 1 day tongue tissue was homogenised to single cells and stained with a live/dead dye, anti-CD45, anti-Ly6G, and anti-CD11b antibody and/or their appropriate isotypes before being analysed by flow cytometry. Flow plots representative of data from 2 independent experiments, gating shown as single CD45^+^ and as percentage (%) of total CD45^+^ cells. (**c**) Weight change of grouped control (dark blue line) and IL-1RAcP^-/-^ mice (light blue line) data from at least 3 independent experiments (n=12 for each group). Each timepoint between groups was compared using an ANOVA with Bonferroni’s post-test. Dotted line represents experimental limit. (**d**) CFU was enumerated from colon, cecum and stomach homogenate at day 4. Each symbol represents one mouse. (**e**) RNA was extracted from tongue and gene expression determined by RT-qPCR. Results were quantified by ΔΔct analysis compared against a naïve sample. Graph displays cumulative data from at least 3 independent experiments (n=6 for each group). Differences between control and IL-1RAcP^-/-^ were analysed by Kruskal-Wallis test with Dunn’s post-test.

**Supplementary figure 5**

(**a-b**) Neutrophils were depleted in control and IL-1RAcP^-/-^ mice by intraperitoneal administration of anti-Ly6G antibody starting 1 day prior to the sublingual administration *of C. albicans ece1*Δ/Δ. (**a**) Weight change of grouped control (dark blue line) and IL-1RAcP^-/-^ mice (light blue line) data from 2 independent experiments (n=8 for each group). Each timepoint was compared to the day 0 value by 2-way ANOVA with Bonferroni’s post-test. (**b**) CFU was enumerated from colon, cecum and stomach homogenate at day 4. Each symbol represents one mouse. (**c**) Neutrophils were depleted in control mice by intraperitoneal administration of anti-Ly6G antibody starting 1 day prior to sublingual administration of *C. albicans* (AHY940). Anti-IL-17A antibody or Anakinra was administered by intraperitoneal administration starting 1 day prior to sublingual administration of *C. albicans* (AHY940). Each line represents grouped control and IL-1RAcP^-/-^ mice data from 2 independent experiments (n=5 for each group). Each timepoint was compared to the day 0 value by 2-way ANOVA with Bonferroni’s post-test. Black dotted line represents experimental limit.

## Notes

Research Support: JRN was supported by grants from the Wellcome Trust (214229_Z_18_Z) and National Institutes of Health (DE022550). SLG was supported by a grant from the National Institutes of Health (DE022550).

### Competing Interest Statement

The authors have declared no competing interest.

