## Supplementary Figures and Table for "The IL-1 Family Controls Acute Mucosal Fungal Infection and Mucosal-Systemic Dissemination"

### Supplementary figure 1

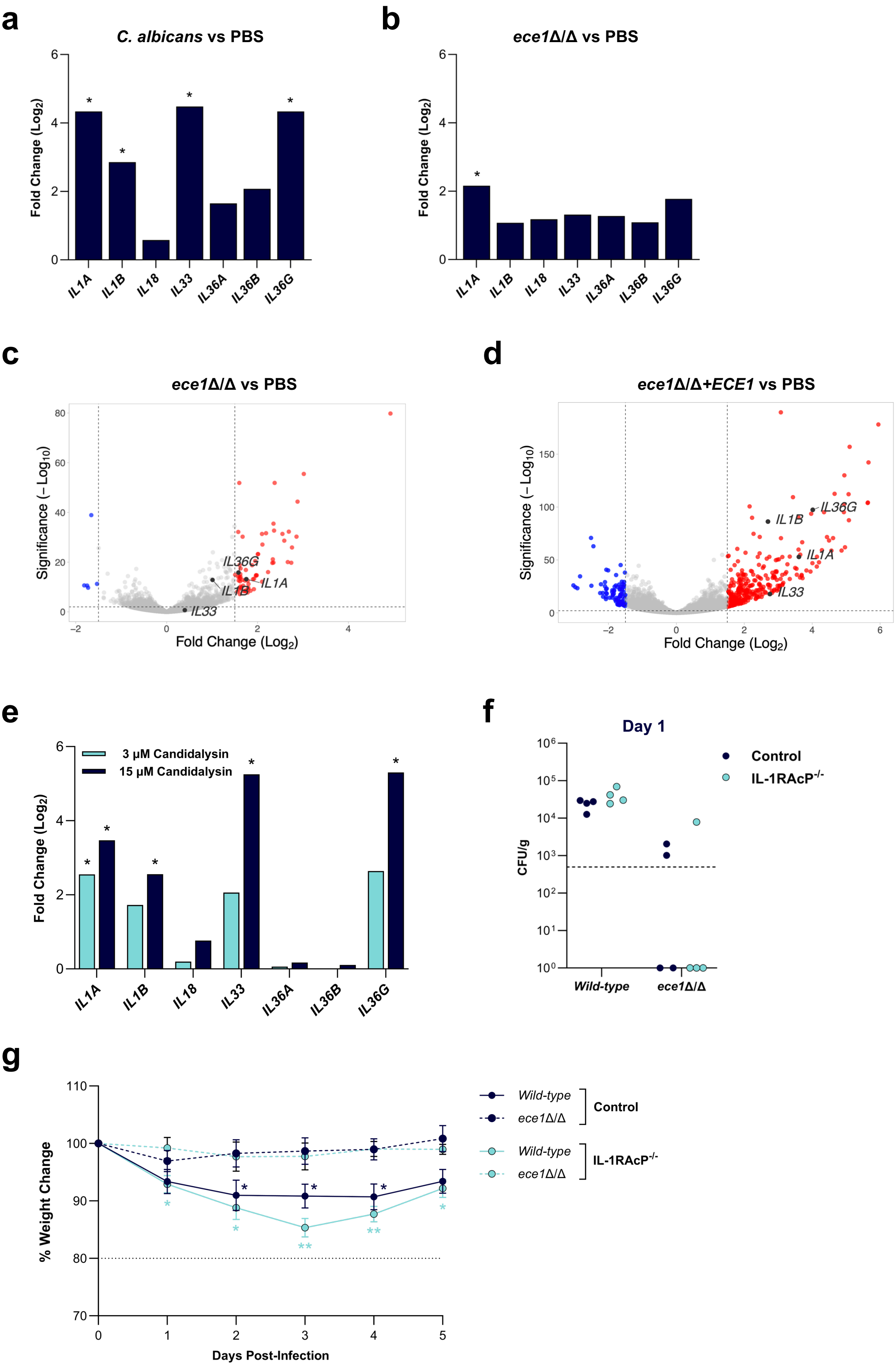

### Supplementary figure 2

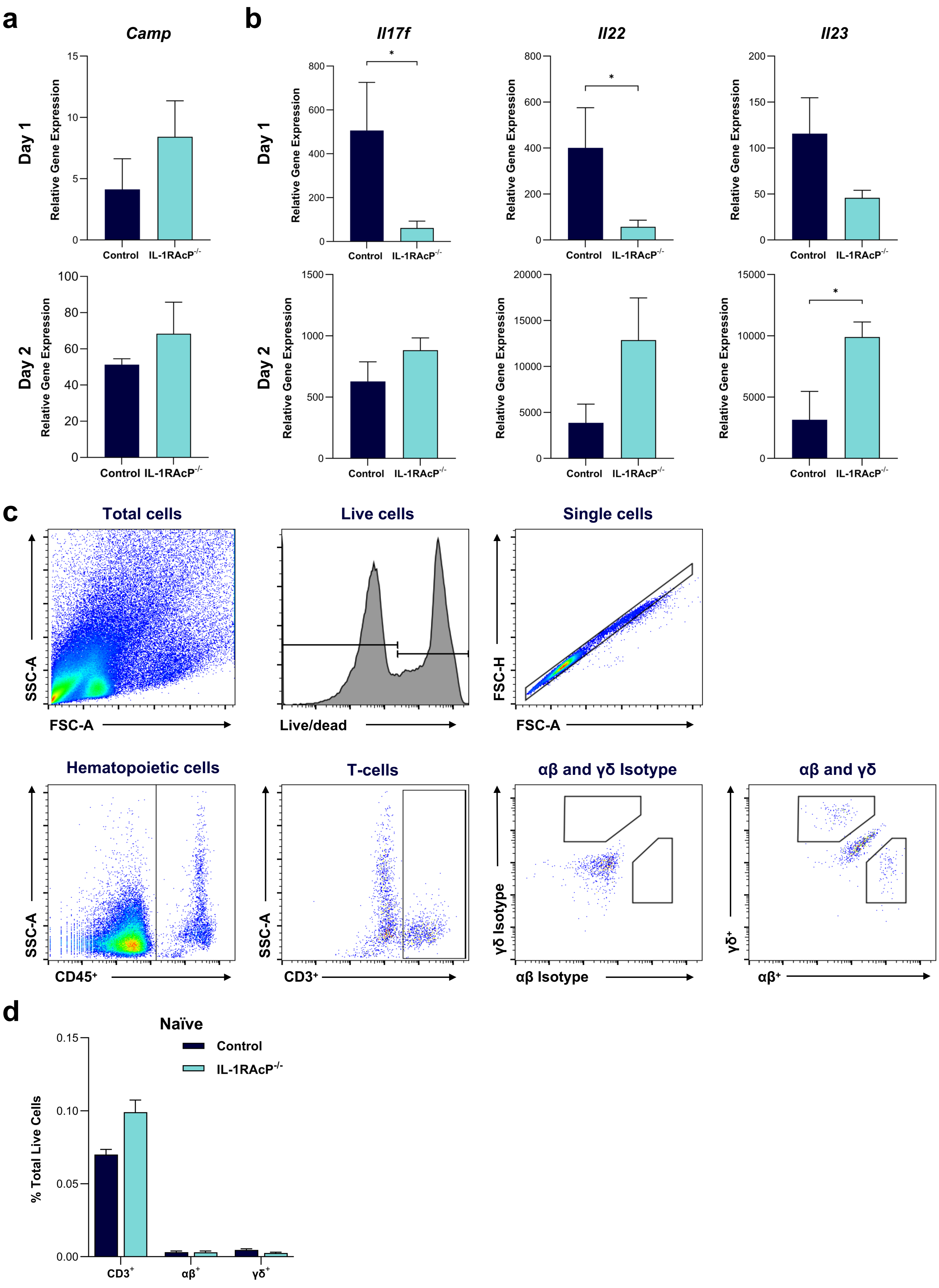

### Supplementary figure 3

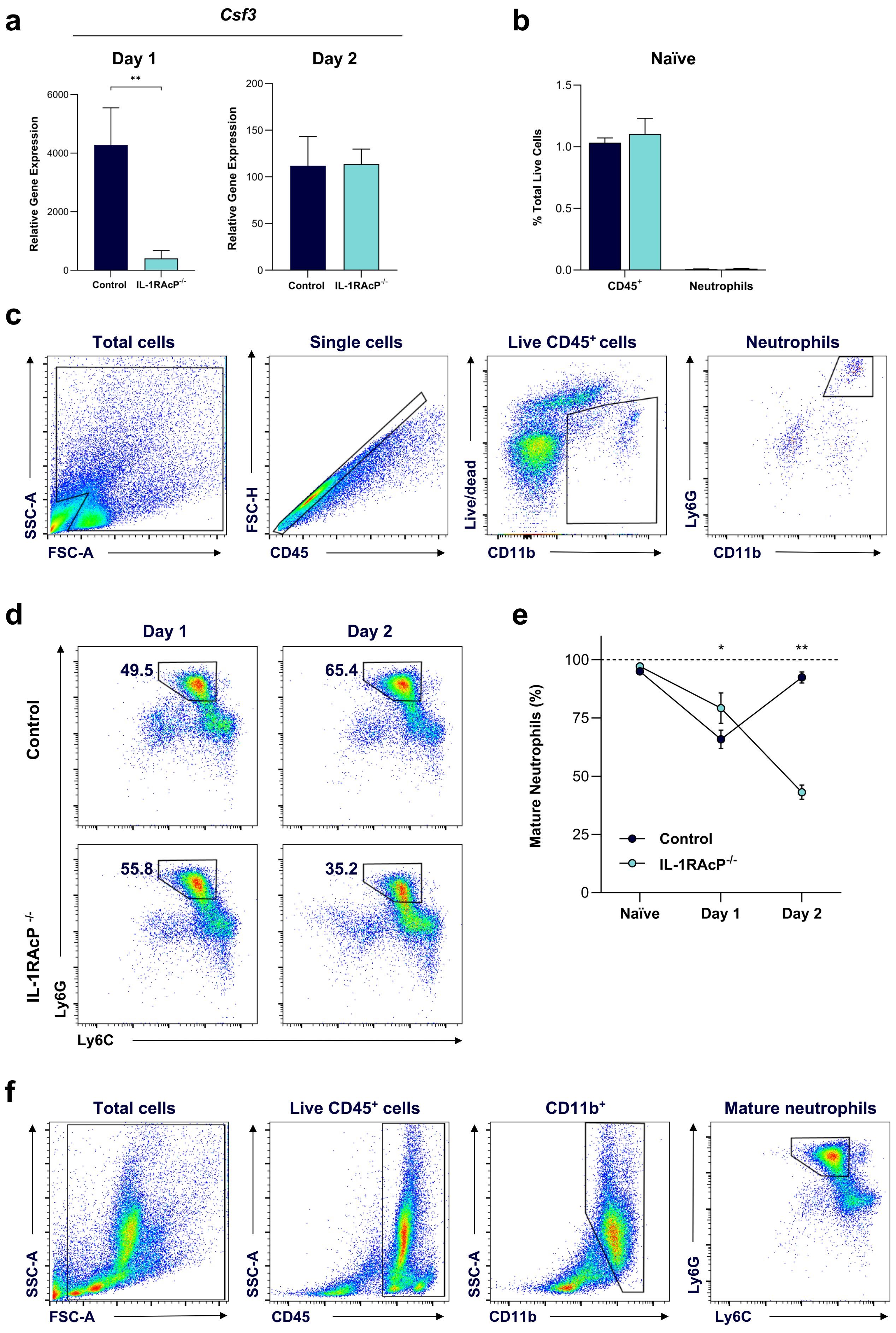

### Supplementary figure 4

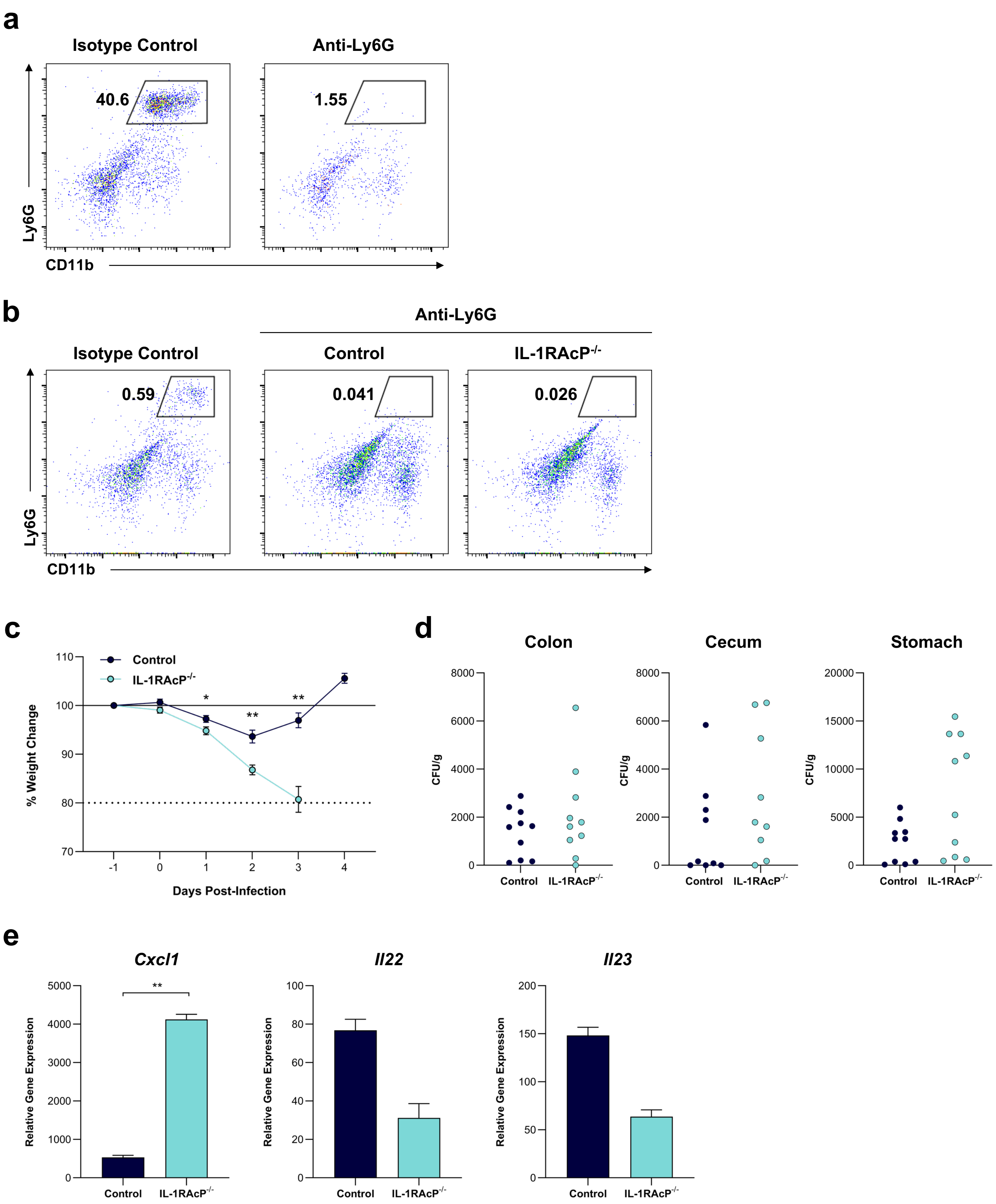

### Supplementary figure 5

**a**

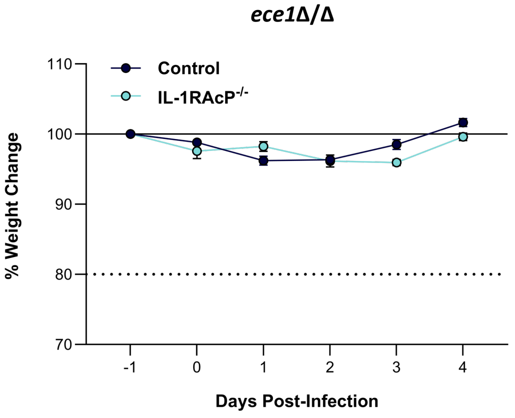


**b**

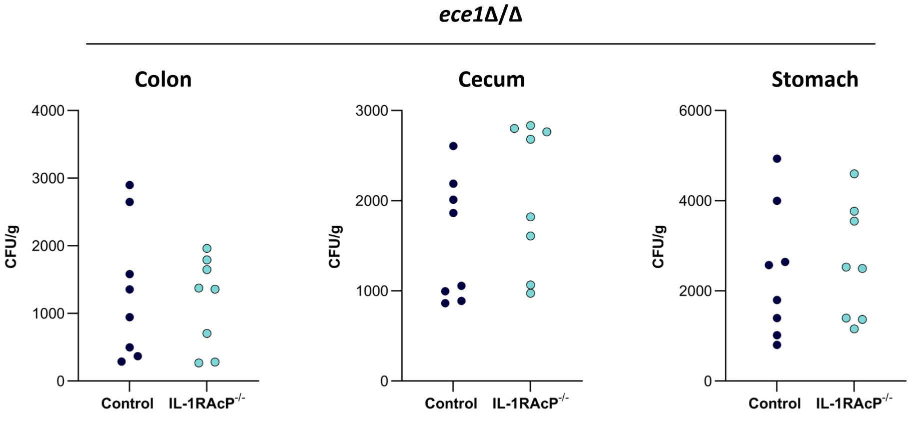


**c**

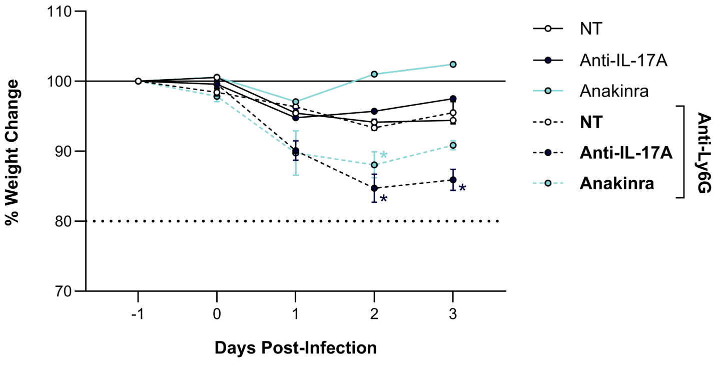

#### Supplementary Table 1

##### RTqPCR Primers

| Gene | Sequences (5’-3’) |
|---|---|
| *Defb3* | F: ACCTTCTGTTTGCATTTCTC |
| | R: GGTCTTCTCTATTTTCTCTTGC |
| *S100a9* | F: AGCCTTGAAGAGCAAGAAG |
| | R: GTCAGGGTGTCCTTCCTTCC |
| *S100a8* | F: ATACAAGGAAATCACCATGC |
| | R: ATATTCTGCACAAACTGAGG |
| *Il17a* | F: CCCCTTTACACCTTCTTTTC |
| | R: ACGTTTCTCAGCAAACTTAC |
| *Camp* | F: AGTGAAGGAGACTGTATGTG |
| | R: ATTTTCTTGAACCGAAAGGG |
| *Il17f* | F: GAAGGCTGGGAACTGTCCTC |
| | R: GGGGTCTCGAGTGATGTTGT |
| *Il22* | F: ATCAGTGCTACCTGATGAAG |
| | R: CATTCTTCTGGATGTTCTGG |
| *Il23* | F: AATAATGCTATGGCTGTTGC |
| | R: CTTAGTAGATTCATATGTCCCG |
| *Cxcl1* | F: AAAGATGCTAAAAGGTGTC |
| | R: GTATAGTGTTGTCAGAAGCC |
| *Csf3* | F: ATGAAGCTAATGGCCCTG |
| | R: CCTGGATCTTCCTCACTTG |

##### Flow Cytometry Antibodies

| Target | Clone | Isotype | Source |
|---|---|---|---|
| Live/Dead (Zombie Green) | | | Biolegend |
| TruStain FcX (anti-mouse CD16/32) | 93 | Rat IgG2a λ | Biolegend |
| Anti-mouse CD45 | 30-F11 | Rat IgG2b κ | Biolegend |
| Anti-mouse CD3 | 17A2 | Rat IgG2b κ | Biolegend |
| Anti-mouse TCRβ | H57-597 | Armenian Hamster IgG | Biolegend |
| Anti-mouse TCRγδ | GL3 | Armenian Hamster IgG | Biolegend |
| Anti-mouse CD11b | M1/70 | Rat IgG2b κ | Biolegend |
| Anti-mouse Ly6G | 1A8 | Rat IgG2a κ | Biolegend |
| Anti-mouse Ly6C | HK1.4 | Rat IgG2c κ | Biolegend |

##### Blocking/Depletion Antibodies

| Target | Clone | Isotype | Source |
|---|---|---|---|
| Vivopure X Anti-Ly6G | 1A8 | Mouse IgG2a | Absolute Antibody |
| InVivoMAb anti-mouse/rat IL-17A | 17F3 | Mouse IgG1 κ | Bio X Cell |
